# Detecting *de novo* mitochondrial mutations in angiosperms with highly divergent evolutionary rates

**DOI:** 10.1101/2020.12.09.418582

**Authors:** Amanda K. Broz, Gus Waneka, Zhiqiang Wu, Matheus Fernandes Gyorfy, Daniel B. Sloan

**Affiliations:** Department of Biology, Colorado State University, Fort Collins, CO 80523; Shenzhen Branch, Guangdong Laboratory for Lingnan Modern Agriculture, Genome Analysis Laboratory of the Ministry of Agriculture, Agricultural Genomics Institute at Shenzhen, Chinese Academy of Agricultural Sciences, 518120 Shenzhen, China

## Abstract

Although plant mitochondrial genomes typically show low rates of sequence evolution, levels of divergence in certain angiosperm lineages suggest anomalously high mitochondrial mutation rates. However, *de novo* mutations have never been directly analyzed in such lineages. Recent advances in high-fidelity DNA sequencing technologies have enabled detection of mitochondrial mutations when still present at low heteroplasmic frequencies. To date, these approaches have only been performed on a single plant species (*Arabidopsis thaliana*). Here, we apply a high-fidelity technique (Duplex Sequencing) to multiple angiosperms from the genus *Silene*, which exhibits extreme heterogeneity in rates of mitochondrial sequence evolution among close relatives. Consistent with phylogenetic evidence, we found that *S. latifolia* maintains low mitochondrial variant frequencies that are comparable to previous measurements in *Arabidopsis. Silene noctiflora* also exhibited low variant frequencies despite high levels of historical sequence divergence, which supports other lines of evidence that this species has reverted to lower mitochondrial mutation rates after a past episode of acceleration. In contrast, *S. conica* showed much higher variant frequencies in mitochondrial (but not in plastid) DNA, consistent with an ongoing bout of elevated mitochondrial mutation rates. Moreover, we found an altered mutational spectrum in *S. conica* heavily biased towards AT➔GC transitions. We also observed an unusually low number of mitochondrial genome copies per cell in *S. conica*, potentially pointing to reduced opportunities for homologous recombination to accurately repair mismatches in this species. Overall, these results suggest that historical fluctuations in mutation rates are driving extreme variation in rates of plant mitochondrial sequence evolution.

## INTRODUCTION

Plant mitochondrial genomes exhibit dramatic variation in rates of nucleotide substitution. Early molecular evolution studies (Wolfe *et al*. 1987; Palmer and Herbon 1988) established that mitochondrial rates in most angiosperms are about an order of magnitude lower than in the nucleus (Drouin *et al*. 2008), which contrasts with the rapid evolution of mitochondrial DNA (mtDNA) in many other eukaryotic lineages (Brown *et al*. 1979; Smith and Keeling 2015; Lavrov and Pett 2016). However, subsequent phylogenetic surveys have identified a number of angiosperms with exceptionally high levels of mtDNA sequence divergence (Palmer *et al*. 2000; Cho *et al*. 2004; Parkinson *et al*. 2005; Mower *et al*. 2007; Sloan *et al*. 2009; Skippington *et al*. 2015; Zervas *et al*. 2019). As such, despite the relatively recent origin and diversification of angiosperms (Barba-Montoya *et al*. 2018), mitochondrial substitution rates are estimated to span a remarkable 5000-fold range across this group (Richardson *et al*. 2013). At one extreme, *Magnolia stellata* has a rate of only ~0.01 synonymous substitutions per site per billion years (SSB), while certain *Plantago* and *Silene* species have estimated rates of >50 SSB (Mower *et al*. 2007; Sloan *et al*. 2012a; Richardson *et al*. 2013). In some cases, rate accelerations appear to be short-lived with bursts of sequence divergence inferred for internal branches on phylogenetic trees followed by reversions to slower rates of sequence evolution (Cho *et al*. 2004; Parkinson *et al*. 2005; Sloan *et al*. 2009; Skippington *et al*. 2017).

The angiosperm genus *Silene* (Caryophyllaceae) is a particularly interesting model for the study of mitochondrial genome evolution and substitution rate variation. This large genus comprises approximately 850 species (Jafari *et al*. 2020) and exhibits rate accelerations that rival the magnitude of other extreme cases in genera such as *Plantago, Pelargonium*, and *Viscum* (Mower *et al*. 2007; Skippington *et al*. 2015). Moreover, *Silene* is distinct because the observed rate accelerations appear to have occurred very recently (<10 Mya), such that close relatives within the genus exhibit radically different substitution rates (Figure 1) (Mower *et al*. 2007; Sloan *et al*. 2009; Rautenberg *et al*. 2012). For example, *S. latifolia* (section *Melandrium)* has retained a substitution rate that is roughly in line with the low levels of sequence divergence found in most angiosperms. In contrast, species such as *S. noctiflora* and *S. conica* represent lineages (sections *Elisanthe* and *Conoimorpha*, respectively) that are within the same subgenus as section *Melandrium* but have highly divergent mtDNA sequence. These large differences among close relatives in *Silene* have enabled comparative approaches to investigate the evolutionary consequences of accelerated substitution rates for mitochondrial genome architecture (Sloan *et al*. 2012a), RNA editing (Sloan *et al*. 2010), mitonuclear coevolution (Sloan *et al*. 2014; Havird *et al*. 2015; Havird *et al*. 2017), and mitochondrial physiology (Havird *et al*. 2019; Weaver *et al*. 2020). Notably, accelerated species such as *S. noctiflora* and *S. conica* also exhibit massively expanded mitochondrial genomes that have been fragmented into dozens of circularly mapping chromosomes (Sloan *et al*. 2012a). However, the mechanisms that cause increased mitochondrial substitution rates in some *Silene* species remain unclear (Havird *et al*. 2017).

**Figure 1.**
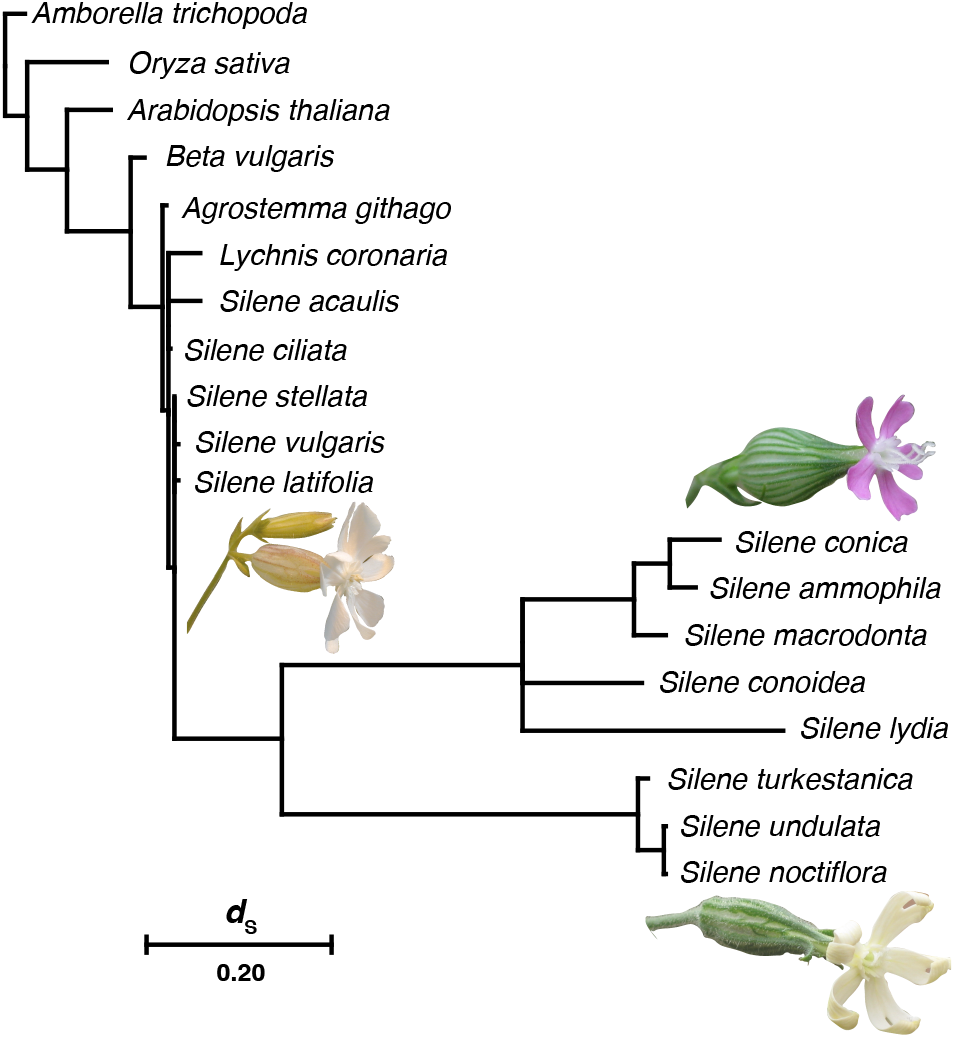
Mitochondrial substitution rate variation among *Silene* species. Branch lengths are scaled based on synonymous substitutions per site (*d*_S_) in a concatenation of three mitochondrial proteincoding genes (*atp1*, *cox3*, and *nad9*) used in previous analyses (Sloan *et al*. 2009; Rautenberg *et al*. 2012). Branch lengths were estimated with PAML v4.9j (Yang 2007), using a constrained topology. Note that the two accelerated clades (section Conoimorpha [represented by *S. conica*] and section Elisanthe [represented by *S. noctiflora*]) were set as sister groups for this analysis, but there is only weak and inconsistent evidence for that relationship (Rautenberg *et al*. 2012; Havird *et al*. 2017; Jafari *et al*. 2020). Images are shown for each of the three focal species in this study (*S. latifolia, S. conica*, and *S. noctiflora*).

Previous studies on *Silene* and other angiosperms have generally concluded that elevated rates of mitochondrial sequence evolution reflect increased mutation rates rather than changes in selection pressure (Cho *et al*. 2004; Parkinson *et al*. 2005; Mower *et al*. 2007; Sloan *et al*. 2009; Sloan *et al*. 2012a; Skippington *et al*. 2015). Increases are especially pronounced at synonymous sites, which are thought to experience very limited effects of selection in plant mtDNA (Sloan and Taylor 2010; Wynn and Christensen 2015). Likewise, the ratio of nonsynonymous to synonymous substitutions (*d*_N_/*d*_S_) is still low in accelerated species (Sloan *et al*. 2012a), indicating that purifying selection on mitochondrial genes remains strong. In *Silene* species with rapidly evolving mtDNA, there does not appear to be a genome-wide increase in synonymous substitution rates in the nucleus or plastids (Rautenberg *et al*. 2012; Sloan *et al*. 2012b), suggesting that the accelerated point mutation rates are largely specific to the mitochondria. A number of mechanisms have been hypothesized to explain cases of increased mitochondrial mutation rates. However, thus far, all inferences about variation in plant mitochondrial divergence are based on long-term patterns of sequence divergence across species rather than direct detection and analysis of *de novo* mutations, making it difficult to investigate the actual role of mutation.

Recent advances in high-fidelity DNA sequencing have improved our ability to distinguish signal from noise and detect *de novo* mutations (Salk *et al*. 2018; Sloan *et al*. 2018). In particular, Duplex Sequencing (Schmitt *et al*. 2012; Kennedy *et al*. 2014) has been used to obtain error rates as low as ~2 × 10^-8^ per bp, allowing for accurate identification of mitochondrial variants that are still present at low heteroplasmic frequencies within tissue samples (Kennedy *et al*. 2013; Ahn *et al*. 2016; Hoekstra *et al*. 2016; Arbeithuber *et al*. 2020; Wu *et al*. 2020a). Duplex Sequencing works by attaching random barcodes to each original DNA fragment and independently sequencing the two complementary DNA strands from that fragment multiple times each to produce a highly accurate double-stranded consensus. To date, *Arabidopsis thaliana* is the only plant system to have been analyzed with this method (Wu *et al*. 2020a). Here, we apply Duplex Sequencing to detect *de novo* mitochondrial and plastid mutations in three *Silene* species previously inferred to have dramatically different histories of mitochondrial mutation.

## MATERIALS AND METHODS

### Plant lines and growth conditions

A single family of full siblings was grown for each of three *Silene* species. Families were taken from lines with previously sequenced mitochondrial and plastid genomes: *S. latifolia* UK2600, *S. noctiflora* OSR, and *S. conica* ABR (Sloan *et al*. 2012a; Sloan *et al*. 2012b). The latter two species are hermaphroditic and readily self-fertilize, so families were derived from selfed parents. In contrast, *S. latifolia* is dioecious and exhibits substantial inbreeding depression from full-sib crosses (Teixeira *et al*. 2009). Therefore, we crossed a female from the UK2600 line with a male from a different line (Kew 32982, obtained from the Kew Gardens Millenium Seed Bank) to produce the full-sib family used in this study. Seeds were germinated on ProMix BX soil mix and grown in a growth room under short-day conditions (10-hr/14-hr light/dark cycle). All three species were grown for approximately seven weeks to produce sufficiently large rosettes for organelle DNA extractions.

### Organelle DNA extractions and Duplex Sequencing

Each full-sib family was divided into three biological replicates, consisting of approximately 30-40 individual plants per replicate. A total of 35 g of rosette tissue was harvested from each replicate and used for simultaneous extraction of enriched mtDNA and cpDNA as described previously (Wu *et al*. 2020a; Wu *et al*. 2020b). Mitochondria were isolated by differential centrifugation followed by DNase I treatment to remove contaminating DNA not protected by intact mitochondrial membranes. Chloroplasts were isolated on discontinuous sucrose gradients. Following DNA extraction, Duplex Sequencing libraries were constructed as described previously (Wu *et al*. 2020a). These libraries were multiplexed and sequenced with 2×150 bp reads on an Illumina NovaSeq 6000 S4 Lane (NovoGene).

### Shotgun Illumina sequencing of total-cellular DNA samples for *k*-mer database construction

Detection of low-frequency mitochondrial heteroplasmies presents a number of technical challenges that can lead to false positives. In particular, plant nuclear genomes harbor numerous insertions of mtDNA and cpDNA fragments (which are known as “numts” and “nupts”, respectively) that can differ in sequence from the mitochondrial or plastid genomes because they accumulate mutations over time (Huang *et al*. 2005; Noutsos *et al*. 2005; Lough *et al*. 2008; Hazkani-Covo *et al*. 2010; Michalovová *et al*. 2013). Therefore, contaminating nuclear DNA in mtDNA and cpDNA samples can mimic low-frequency *de novo* mitochondrial and plastid mutations. Because numts and nupts often closely resemble the mitochondrial and plastid genome sequence, they can be problematic to accurately assemble in nuclear genome sequencing projects even in high quality reference assemblies (Stupar *et al*. 2001). As such, it can be difficult to identify and filter out numt- and nupt-derived variants based only on reference genome assemblies. We have found that an effective alternative is to use raw reads from total-cellular sequencing datasets to generate a database of counts of all sequences of a specified length *k*, which are referred to as *k*-mers (Wu *et al*. 2020a).

The premise of this approach is that variants associated with numts or nupts should be abundant in total-cellular sequencing datasets (as quantified by counting corresponding *k*-mers in the raw reads) and match the level expected for other nuclear sequences. However, for reasons discussed below (see Results), we expect this filtering approach to be more reliable for numts than for nupts.

To generate a *k*-mer database for each species we extracted total-cellular DNA from mature leaf tissue using the Qiagen DNeasy kit. For *S. noctiflora* and *S. conica*, the sampled individuals were from the same inbred line as the full-sib family used for Duplex Sequencing but separated by at least three generations. In contrast to the inbred history of the *S. noctiflora* and *S. conica* lines, *S. latifolia* is expected to be highly heterozygous, including for numt and nupt alleles. Therefore, in order to capture all numt and nupt alleles that might be segregating in the *S. latifolia* full-sib family, we generated total-cellular samples for both parents that were crossed to generate the family. Sampling the male parent in this *S. latifolia* cross also provides reads from its actual mitochondrial genome sequence, which can identify potential false positives resulting from low-frequency paternal transmission of mtDNA. Such “paternal leakage” has been identified in outcrossing *Silene* species (McCauley *et al*. 2005; Bentley *et al*. 2010).

Illumina libraries were generated from each total-cellular DNA sample using the New England Biolabs NEB FS II Ultra kit with approximately 100 ng of input DNA, a 20 min fragmentation time, and eight cycles of PCR. The *S. latifolia* and *S. noctiflora* libraries were multiplexed and sequenced on the same NovaSeq 6000 lane as the above Duplex Sequencing libraries. The *S. conica* library was sequenced separately with 2×150 bp reads on an Illumina HiSeq 4000 lane. The resulting raw reads were used to generate databases of *k*-mer counts (*k* = 39 bp) with KMC v3.0.0 (Kokot *et al*. 2017).

### Duplex Sequencing data analysis and variant calling

Duplex Sequencing datasets were processed with a previously published pipeline (https://github.com/dbsloan/duplexseq) (Wu *et al*. 2020a). This pipeline uses Duplex Sequencing barcodes (i.e., unique molecular identifiers) to group raw reads into families corresponding to the two complementary strands of an original DNA fragment, requiring a minimum of three reads for each strand. After trimming barcodes and linker sequences, the consensus sequences for each double-stranded family were mapped to the reference mitochondrial and plastid genomes for the corresponding species. Because of known sequencing artefacts associated with end repair and adapter ligation (Kennedy *et al*. 2014), the 10 bp at the end of each read were excluded when identifying variants and calculating sequencing coverage.

Reads that contained a single mismatch relative to the reference sequences were used to identify single nucleotide variants (SNVs). Variants with a *k-mer* count of 10 or greater in the corresponding *k*-mer database were excluded as likely numts or nupts. This *k*-mer filtering also ensured elimination of false positives due to paternal leakage from the *S. latifolia* male parent or errors in the published reference sequence. The latter is an important concern because the published *S. conica* mitochondrial genome sequence is a draft assembly due to its extreme repetitiveness (Sloan *et al*. 2012a). Accordingly, we also excluded variants if the corresponding reference sequence was not detected in the *k*-mer database to account for sites with ambiguities (Ns) or possible errors in the reference. Variants were also filtered using the pipeline’s --recomb_check option, which identifies SNVs that can be explained by recombination between nonidentical repeat sequences within the mitochondrial genome rather than *de novo* point mutations (Davila *et al*. 2011). Finally, the pipeline’s --contam_check option was used to provide the reference genome sequences from the other *Silene* species for filtering of variants arising from contamination between multiplexed libraries.

To calculate SNV frequencies for each library, the number of reads containing an SNV (after filtering) was divided by the total bp of duplex consensus sequence mapped to the genome. Differences in SNV frequency among species were tested with a one-way ANOVA implemented, using the aov function in R v3.6.3 followed by *post hoc* pairwise comparisons with the TukeyHSD function.

For calling mitochondrial SNVs, we were able to use the cpDNA Duplex Sequencing libraries to increase our mitochondrial genome coverage because they contained a substantial amount of contaminating mtDNA reads resulting from incomplete enrichment of cpDNA. Importantly, being able to use mitochondrial reads from the cpDNA library for *S. conica* biological replicate 2 was key because sequencing of the mtDNA library for that replicate failed (see Results). However, we did not do the reverse (use mtDNA libraries to supplement plastid datasets) because the mtDNA libraries were expected to have a far higher ratio of nuclear to plastid DNA than the cpDNA libraries, exacerbating the challenges associated with filtering nupts.

### Analysis of organelle genome copy number

To estimate the copy number of mitochondrial and plastid genomes based on the total-cellular sequencing datasets, raw reads were trimmed with cutadapt v1.16 (Martin 2011) to remove adapter sequences with an error tolerance of 0.15 and to trim low-quality ends with a q20 threshold. Read pairs with a minimum length of 50 bp each after trimming were retained. Trimmed reads were then mapped to reference mitochondrial and plastid genomes for the corresponding species using bowtie v2.2.23 (Langmead and Salzberg 2012), and position-specific coverage data were extracted from the resulting alignment (BAM/SAM) files. Average coverage was summarized for 2-kb windows tiled across each organelle genome. To estimate the number of mitochondrial and plastid genomes per nuclear genome copy, we assumed that the remaining unmapped reads were all nuclear. We used the total length of these unmapped reads divided by the nuclear genome size of the corresponding species to estimate the average nuclear genome coverage and obtain ratios of organelle to nuclear coverage. Nuclear genome size estimates of 2.67, 2.78, and 0.93 Gb were used for *S. latifolia, S. noctiflora*, and *S. conica*, respectively (Williams *et al*. 2020).

The above analysis of genome copy number identified a surprisingly low number of mitochondrial genome copies in *S. conica*. To validate this unexpected result and compare stoichiometry of mitochondrial, plastid and nuclear genomes across multiple tissue samples in *S. conica*, we performed a droplet digital PCR (ddPCR) analysis. We used the same *S. conica* total-cellular DNA extraction that was used for the Illumina shotgun sequencing. In addition, we harvested leaf tissue from three individuals from a different batch of *S. conica* ABR plants. For both the original sample and the newer samples, tissue was harvested from the largest rosette leaves. The original sample was grown in a growth chamber under long-day conditions (16-hr/8-hr light/dark cycle) and harvested after 28 days, whereas the newer samples were grown on light racks in a growth room under short-day conditions (10-hr/14-hr light/dark cycle) and harvested after 40 days. The tissue sampling also differed in that the entire rosette leaf was sampled for the newer replicates, whereas only the distal half of the leaf was taken for the original DNA extraction. In both cases, DNA extractions were performed with a Qiagen DNeasy Kit. A total of six ddPCR markers were developed, with two each targeting the mitochondrial, plastid and, nuclear genomes (Table S1). Each ddPCR reaction was set up in a 20 μl volume, containing Bio-Rad QX200 ddPCR EvaGreen Supermix and 0.2 μM of each primer. For mitochondrial and nuclear markers, 5 ng of total-cellular DNA was used as template. To avoid saturation with the higher-copy-number plastid makers, a 200-fold dilution (25 pg) of the template DNA was used. The reaction volumes were then emulsified in Bio-Rad QX200 Droplet Generation Oil for EvaGreen, using the Bio-Rad QX200 Droplet Generator. PCR amplification was performed on a Bio-Rad C1000 Touch thermal cycler with an initial incubation at 95 °C for 5 min, 40 cycles of 90 °C for 30 s and 60 °C for 1 min, followed by a 4 °C incubation for 5 min and a final 90 °C incubation for 5 min before holding at 4 °C. After amplification, droplets were analyzed on a Bio-Rad QX200 Droplet Reader, and the absolute copy number of each PCR target was estimated based on a Poisson distribution in the Bio-Rad QuantaSoft package. Mitochondrial:nuclear and plastid:nuclear ratios were calculated by dividing organellar marker copy numbers by the mean of the estimates for the two nuclear markers for a given sample.

### Data Availability

All duplex sequencing and shotgun Illumina sequencing reads were deposited to the NCBI Sequence Read Archive (SRA) under BioProject PRJNA682809 (Table S2).

## RESULTS

### Duplex Sequencing yield and read mapping

Each of the Duplex Sequencing libraries produced between 88 M and 142 M read pairs (Table S2), with the exception of the *S. conica* mtDNA library for biological replicate 2, which was not sequenced due to an apparent loading error. Nevertheless, we were still able to calculate mitochondrial SNV frequencies for *S. conica* replicate 2 by taking advantage of contaminating mitochondrial reads in the enriched cpDNA library for that replicate. The large number of reads per library translated into sizeable single-stranded read families for construction double-stranded consensus sequences, with modal values of >10 reads per family for most libraries (Figure S1). The three species differed in their degree of enrichment in the mtDNA libraries. For both *S. noctiflora* and *S. conica*, an average of 79% of the sequences in the mtDNA libraries could be mapped to the reference mitochondrial genome, whereas only 30% of the sequences in the *S. latifolia* mitochondrial libraries mapped, indicating a much lower level of mtDNA enrichment (Table S2). For all three species, between 64% and 67% of sequences from the cpDNA libraries mapped to the plastid genome, with a substantial number of contaminating sequences mapping to the mitochondrial genome (8%, 27%, and 31% for *S. latifolia, S. noctiflora*, and *S. conica*, respectively). To make use of as much mitochondrial data as possible, we combined all mitochondrial-derived sequences from both sample types. After collapsing raw reads into duplex consensus sequences and mapping them to the reference genome, we obtained between 85 and 382 Mb of mapped mitochondrial sequence data for each replicate (based on combing coverage from both mtDNA and cpDNA libraries; Table S2). For plastid genome coverage, we relied solely on the cpDNA libraries, which yielded between 203 and 319 Mb of coverage per replicate (Table S2).

### Three *Silene* species differ in their frequencies of mitochondrial SNVs but show little variation in plastid SNV frequencies

Using the variant calls from the duplex consensus sequences (File S1), we calculated the frequency of mitochondrial SNVs per mapped bp for each *Silene* replicate and compared those values to previously published estimates from *A. thaliana* (Wu *et al*. 2020a). After applying filtering criteria to exclude false positives associated with numts, low-frequency paternal transmission, chimeric recombination products, or errors in the reference sequence (see Materials and Methods), we found significant variation in (log-transformed) mitochondrial SNV frequency among species (one-way ANOVA, *p* = 5.3 × 10^-6^; Figure 2, Table S3).

**Figure 2.**
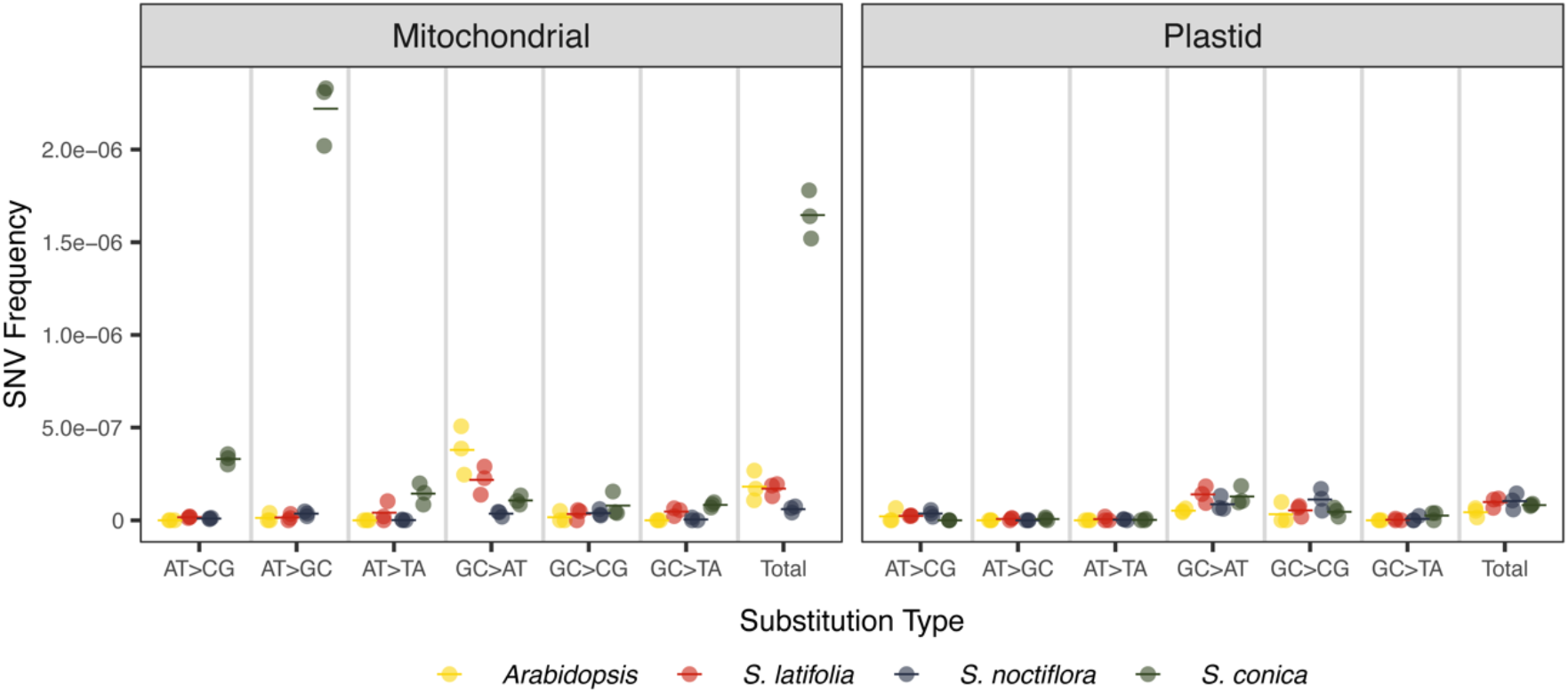
Variation in mitochondrial and plastid SNV frequencies and spectra across *Arabidopsis* and *Silene* species. SNV frequencies are calculated as the total number of observed singlenucleotide mismatches in duplex consensus sequence datasets dividided by the total base-pairs of mitochondrial or plastid genome coverage in those datasets (i.e., GC coverage, AT coverage, or total coverage depending on the SNV type). Three biological replicates (circles) are shown for each species, with the mean of those replicates indicated with a horizontal line. The *Arabidopsis* data points are taken from Wu et al. (2020). The same data are plotted on a log scale in Figure S2.

The three biological replicates of *S. latifolia* had a mean mitochondrial SNV frequency of 1.7 × 10^-7^ per bp. *Silene latifolia* was selected for this study because it exhibited very little mitochondrial rate acceleration in previous phylogenetic analyses, suggesting that it has retained the slow rate of sequence evolution that is characteristic of most plant mitochondrial genomes (Mower *et al*. 2007; Sloan *et al*. 2009). Accordingly, the *S. latifolia* estimate was very similar to our previous estimate of 1.8 × 10^-7^ per bp for the mitochondrial SNV frequency in wild type *A. thaliana* Col-0 (Wu *et al*. 2020a), another plant species with a typically low rate of mitochondrial sequence evolution (Mower *et al*. 2007). In contrast to the low historical substitution rates in *S. latifolia* and *Arabidopsis, S. noctiflora* exhibits highly accelerated mitochondrial sequence evolution since diverging from other major lineages within the genus *Silene* (Figure 1). However, we did not find an elevated frequency of mitochondrial SNVs in our Duplex Sequencing analysis of *S. noctiflora*, suggesting that this species may have experienced a reversion to lower mutation rates. In fact, the mean SNV frequency in *S. noctiflora* was 6.1 × 10^-8^ per bp, which was approximately 3-fold lower than in *S. latifolia* or *A. thaliana* (Tukey’s HSD *post hoc* test, *p* = 0.01 for both comparisons). The most noteworthy variation among species was based on observed SNV frequencies in *S. conica*, representing another *Silene* lineage with a history of rapid mitochondrial sequence divergence (Figure 1). The mean mitochondrial SNV frequency in *S. conica* was 1.7 × 10^-6^ per bp, which was 9-fold to 27-fold higher than in *A. thaliana* and the other two *Silene* species (Tukey’s HSD post hoc test, *p* < 0.0001 for all three comparisons). All of these SNV frequencies substantially exceed the noise threshold of ~2 × 10^-8^ that we previously estimated for this protocol using *Escherichia coli* samples derived from single colonies as “negative controls” (Wu *et al*. 2020a).

*Silene conica* was also distinct in that a large proportion of the identified mitochondrial SNVs (31.7%) were found in two or more biological replicates. Because our biological replicates represented sets of individuals from the same full-sib family, variants that are shared among replicates likely indicate SNVs that were already heteroplasmic in the parent and then inherited by the offspring. In contrast, none of the identified mitochondrial SNVs in either *S. latifolia* or *S. noctiflora* were present in multiple biological replicates. There is reason to expect that our *k*-mer filtering may have introduced bias against detection of inherited heteroplasmies in *S. latifolia* (see Discussion). Nevertheless, even without this filtering, only 3.7% of *S. latifolia* SNVs (and only 1.8% of *S. noctiflora* SNVs) would be present in two or more libraries. Therefore, the filtering does not appear to explain this observed difference between *S. conica* and the other *Silene* species.

Unlike in the mitochondrial genome, we found no evidence that *S. conica* has an elevated frequency of SNVs in its plastid genome, as the three *Silene* species all exhibited similarly low plastid SNV frequencies (Figure 2). We recommend that the estimates of the *Silene* plastid SNV frequencies and spectra be interpreted cautiously because of the nuclear contamination in these libraries and the fact that nupts are more difficult to reliably filter with our *k*-mer database than numts. The challenge that nupts pose relates to the high coverage of true plastid DNA in our total-cellular libraries (>10,000×). As such, even rare sequencing errors in total-cellular libraries have the potential to occur repeatedly and exhibit sizeable counts in our *k*-mer database, which could lead to exclusion of variant calls that are actually true *de novo* mutations. Nevertheless, we can confidently conclude that *S. conica* does not exhibit a major increase in plastid SNV frequency. Even if we performed no *k*-mer filtering whatsoever on the *S. conica* plastid samples, SNV frequencies would only increase by 55% on average (Table S4), leaving them at a level that is still similar to that of the other *Silene* species and more than an order of magnitude lower than the (filtered) mitochondrial SNV frequencies in *S. conica* (Figure 2).

### Variation in mitochondrial mutation spectra among *Silene* species and extreme GC bias in *S. conica*

Previous analysis of low-frequency mitochondrial SNVs in rosette tissue from wild type *A. thaliana* Col-0 (Wu *et al*. 2020a) identified a mutation spectrum dominated by GC➔AT transitions (Figure 2). The slowly evolving *S. latifolia* mitochondrial genome exhibited a bias in this same direction with 57% of all observed SNVs being GC➔AT transitions (Table S3). In contrast, *S. noctiflora* did not show a similar bias. The low overall SNV frequency in *S. noctiflora* makes it difficult to precisely estimate the mutation spectrum, but no single substitution type dominated, as GC➔AT transitions, AT➔GC transitions, and GC➔CG transversions all had similar frequencies in the observed set of SNVs (Figure S2, Table S3). Once again, *S. conica* exhibited the most extreme departure from the other species. Notably, the high rate in *S. conica* was not driven by an increased frequency of the GC➔AT transitions that dominate the spectrum of *A. thaliana* and *S. latifolia*. In fact, the frequency of GC➔AT transitions in *S. conica* was lower than in either of those species. Instead, the high overall SNV frequency was largely the result of a massive increase in the frequency AT➔GC transitions, which account for 77% of the observed SNVs in *S. conica* (Figure 2; Table S3). This species also exhibited a substantial increase in the frequency of AT➔CG transversions (11% of observed SNVs). As such, both of the dominant types of substitutions in the *S. conica* mitochondrial mutation spectrum increase GC content, which is unusual because mutation spectra are generally AT-biased (Hershberg and Petrov 2010; Hildebrand *et al*. 2010; Sloan and Wu 2014).

### Unusually low mitochondrial genome copy number in *Silene conica* rosette tissue

By performing deep sequencing of total-cellular DNA to generate a *k*-mer database for variant filtering, we were also able to estimate the relative copy number of mitochondrial, plastid, and nuclear genomes in each of the three *Silene* species (Figures 3 and S3–S5, Table S5). We found similar plastid genome copy numbers across species, with mean estimates of 378, 263, and 275 plastid genome copies per nuclear genome copy for *S. latifolia*, *S. noctiflora*, and *S. conica*, respectively. If we assume that each cell is diploid and has not yet undergone DNA replication, the plastid genome copy numbers per cell would be double those values, but that may be an underestimate because many plants undergo extensive endoreduplication, in which the nuclear genome replicates without subsequent cell divisions, resulting in cells with higher nuclear ploidies (Joubes and Chevalier 2000). The number of mitochondrial genome copies was surprisingly low in *S. conica*, with an average of only 0.38 copies per nuclear genome copy. In contrast, *S. latifolia* and *S. noctiflora* had an average of 47.7 and 9.7 mitochondrial genome copies per nuclear genome copy, respectively, which is more consistent with estimates from other plants (Preuten *et al*. 2010; Oldenburg *et al*. 2013; Shen *et al*. 2019).

**Figure 3.**
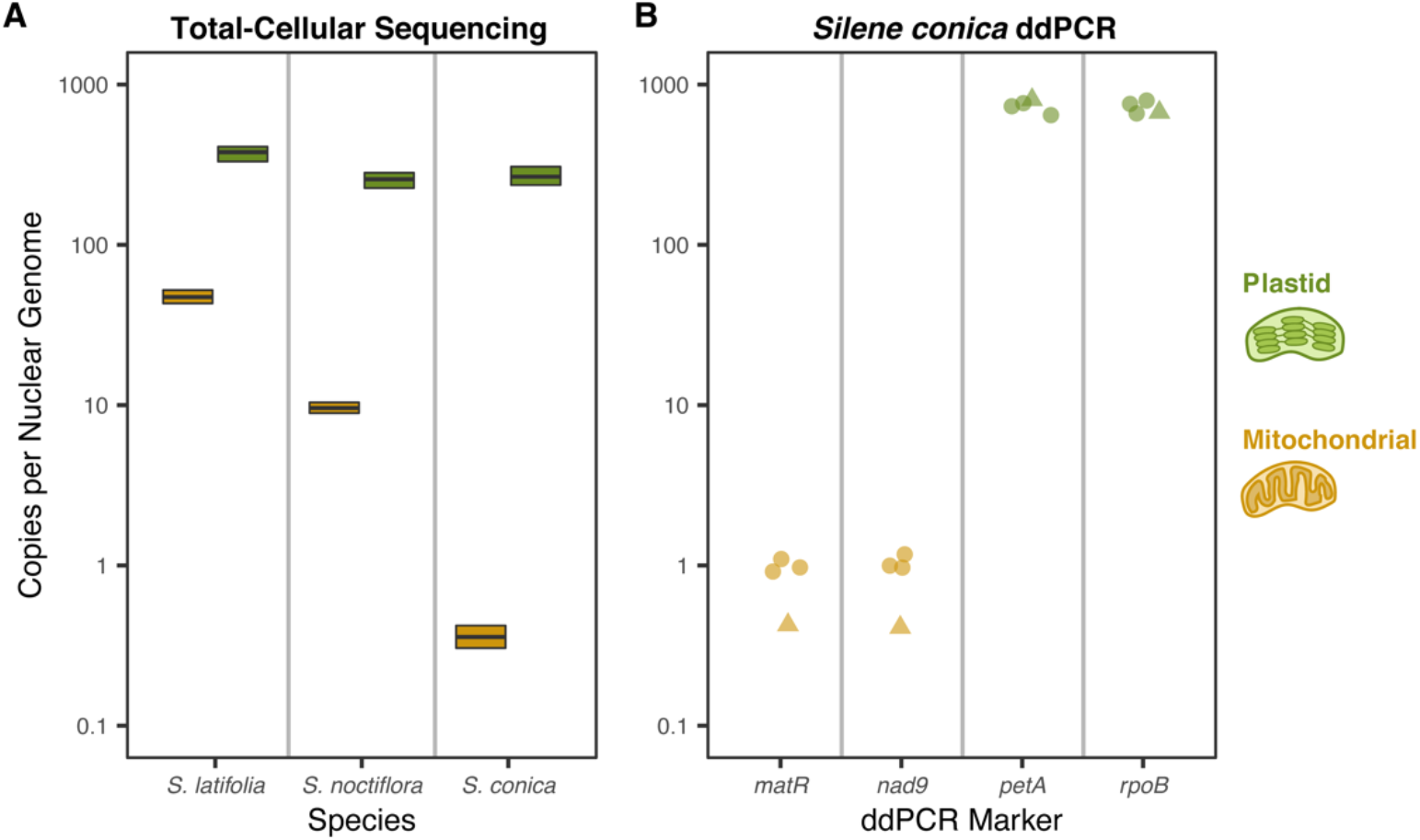
Variation in mitochondrial genome copy number among *Silene* species. (A) Average mitochondrial, plastid, and nuclear genome copy number were estimated from total-cellular Illuimina shotgun sequencing of leaf tissue from each species. Boxplots show median and interquartile ranges for the ratio of organelle genome copy number to nuclear genome copy number based on scanning the organelle genomes in 2-kb windows. Green and gold boxplots correspond to plastid:nuclear and mitochondrial:nuclear ratios, respectively. (B) ddPCR analysis of mitochondrial and plastid genome copies per nuclear genome copy in *S. conica*. Points are shown for two mitochondrial markers (*matR* and *nad9*) and two plastid markers (*petA* and *rpoB*). Estimates for copy number ratios were generated by dividing each mitochondrial or plastid value by the average copy number of two nuclear markers for the corresponding sample (Table S1). The triangles indicate ddPCR estimates for the sample taken from the same DNA extraction used in the original sequencing analysis in part (A). The circles represent the three new samples collected for this ddPCR analysis.

To validate the finding of extremely low mitochondrial genome copy number in *S. conica*, we performed a ddPCR analysis with two markers each for the mitochondrial, plastid, and nuclear genomes. We first analyzed DNA from the same extraction that was originally used for the total-cellular Illumina shotgun sequencing, obtaining an estimate of 0.42 mitochondrial genomes per nuclear genome, in close agreement with our estimate of 0.38 from the sequencing data. Based on this ddPCR analysis, we also estimated that there were 737 copies of the plastid genome per nuclear genome copy, which was 2.7-fold higher than our original estimate (Figure 3), possibly indicating that our sequencing estimate was downwardly biased for plastid genome copy number. We then analyzed leaf DNA collected from three new *S. conica* plants that were grown separately from the originally sampled plant to assess whether the original DNA extraction was anomalous in some way. These three new samples also produced extremely low estimates for the number of mitochondrial genomes copies with a mean of 1.02 per nuclear genome copy (Figure 3 and Table S6). Therefore, these new *S. conica* samples exhibited a small increase in the mitochondrial:nuclear ratio relative to our original sample but still fell well below typical observations for plant cells.

## DISCUSSION

### The challenges of detecting *de novo* mutations in plant organelle genomes

High-fidelity techniques such as Duplex Sequencing (Schmitt *et al*. 2012) have been key innovations to address the challenge of detecting and quantifying rare mutations (Salk *et al*. 2018; Sloan *et al*. 2018), but some sources of false positives remain. The potential misidentification of numts and nupts as *de novo* mutations was a particular concern in this study. High quality nuclear genome assemblies are not available for *Silene*, so it is not possible to use a reference genome to identify and filter numt- and nupt-associated variants. Moreover, our mitochondrial and plastid DNA preparations only reached moderate levels of enrichment, leaving substantial amounts of contaminating nuclear DNA (Table S2). Our approach to avoid erroneous numt and nupt variant calls was based on filtering using a *k*-mer database derived from total-cellular sequencing (see Materials and Methods), but there are some concerns about balancing false positives and false negatives that should be considered.

In particular, there is a risk that *k*-mer filtering may exclude true heteroplasmies if they are shared between the total-cellular sample used to generate the *k*-mer database and the family used for Duplex Sequencing. We reduced the risk of this in *S. conica* and *S. noctiflora* by using individuals separated by at least three generations for constructing the *k*-mer database. Therefore, low-frequency heteroplasmies would have to have been maintained across multiple generations to lead to improper exclusion of true mitochondrial variants. Although transmission of heteroplasmies across generations certainly occurs (McCauley 2013; Zhang *et al*. 2018; Mandel *et al*. 2020), the segregational loss of low-frequency variants should greatly reduce the magnitude of this problem. In contrast, for the outcrossing species *S. latifolia*, we used both parents of the full-sib family to construct total-cellular *k*-mer databases in order to screen for possible numts and nupts in all contributing nuclear haplotypes. As such, variants that were heteroplasmic in the *S. latifolia* mother and transmitted to her offspring might have been improperly filtered because of their presence in the total-cellular *k*-mer database, resulting in a potential downward bias on our estimate of the overall frequency of SNVs in *S. latifolia*.

Despite the uncertainty that this introduces into SNV frequency estimates, we feel that the major qualitative conclusions of this study are robust to the challenges of numt and nupt artefacts. For example, the finding that *S. noctiflora* appears to have “reverted” to a low SNV frequency is not sensitive to *k*-mer filtering. Even if no *k*-mer filtering whatsoever were performed for *S. noctiflora*, it would still exhibit an SNV frequency lower than the filtered values from the other three species (Table S3). Likewise, even if we did not perform any numt filtering on the *S. latifolia* SNVs (which would almost certainly lead to a substantial overestimation of true mitochondrial mutations), the SNV frequency for *S. latifolia* would still not reach the highly elevated levels in *S. conica*. Therefore, the key distinctions among the three species in mitochondrial SNV frequency appear to hold even though some caution is warranted in interpreting the specific frequency estimate in *S. latifolia*. Furthermore, as noted in the Results, the conclusion that plastid SNV frequencies remain low in *S. conica* is not sensitive to *k*-mer filtering, as removing this filtering step only produces a modest increase in the estimate for *S. conica* plastid mutations (Table S4).

### Heteroplasmic SNV frequencies support a history of mitochondrial mutation rate acceleration and reversion in *Silene*

The high frequency of mitochondrial SNVs captured by Duplex Sequencing in *S. conica* tissue (Figure 2) is consistent with previous interpretations that increased mutation rates are driving accelerated mitochondrial genome evolution in this and other atypical plant species (Cho *et al*. 2004; Parkinson *et al*. 2005; Mower *et al*. 2007; Sloan *et al*. 2009; Sloan *et al*. 2012a; Skippington *et al*. 2015). This view has become the consensus because accelerations are evident at relatively neutral positions like synonymous sites over phylogenetic timescales, but *de novo* mitochondrial mutations have never been directly investigated in these high-rate plant lineages until now. The huge increase in AT➔GC transitions that dominated the *S. conica* mutation spectrum (Figure 2) coincides with the most common type of misincorporation observed in steady-state kinetic analysis of the *Arabidopsis* mitochondrial DNA polymerases, which appear to be prone to misincorporating Gs opposite Ts (Ayala-García *et al*. 2018). Therefore, it is possible that the increased mitochondrial substitution rate and biased mutation spectrum in *S. conica* reflect a reduced ability to repair mismatches created by polymerase misincorporations. A disproportionate increase in AT➔GC transitions was also observed in *Arabidopsis* mutants lacking a functional copy of *MSH1* (Wu *et al*. 2020a), a gene that may be involved in repair of mismatches via homologous recombination (Christensen 2014; Wynn *et al*. 2020). An intact and transcribed copy of *MSH1* is retained in *S. conica* (Havird et al. 2017), but its function and expression patterns have not been investigated. Alternatively, GC-biased gene conversion has also been hypothesized as a mechanism to explain skewed substitution patterns in some plant mitochondrial genomes (Liu *et al*. 2020).

The extreme bias towards AT➔GC transitions in *S. conica* mitochondrial SNVs (Figure 2) is not entirely consistent with longer-term patterns of mitochondrial sequence divergence in this species. The genome-wide GC content in *S. conica* (43.1%) is only slightly higher than in congeners like *S. latifolia* (42.6%), *S. noctiflora* (40.8%), and *S. vulgaris* (41.8%) (Sloan *et al*. 2012a). An analysis that used plastid DNA insertions in mitochondrial genomes as neutral markers did find that *S. conica* had unusually high transition:transversion ratio compared to other angiosperms (Sloan and Wu 2014), which is consistent with the Duplex Sequencing results. However, it did not detect the strong GC bias that we observed in the current study.

These discrepancies between phylogenetic patterns and our duplex sequencing data raise two obvious possibilities. First, the observed SNVs in this study may not reflect the spectrum of heritable mutations because they are measured from rosette tissue and thus are expected to include some *de novo* mutations that occurred in vegetative tissues and were not inherited from the mother or transmitted to future generations. Our choice to sample whole rosettes (as opposed to more targeted “germline” tissue such as dissected meristems) reflected the practical need to obtain sufficient quantities of mtDNA and cpDNA for Duplex Sequencing library construction. Although it is important to recognize the observed SNVs include some mutations that are not heritable, we do not believe that this is likely to be the primary explanation for inferred differences in mitochondrial mutation spectra. A large proportion of the *S. conica* SNVs were shared across more than one biological replicate, implying that they were inherited from a heteroplasmic mother and thus transmitted across generations. Furthermore, these shared variants were even more skewed towards AT➔GC transitions than variants that were only detected in a single replicate (Table S7), suggesting that heritable mutations do indeed exhibit a very strong GC bias. Relatedly, the fact that *S. conica* had such a large number of shared SNVs compared to the other two *Silene* species (see Results and Table S7) supports the conclusion that the higher overall SNV frequency in *S. conica* is not solely due to a higher mutation rate in vegetative tissue and indeed reflects an elevated rate of heritable mutations.

Second, it is possible that the mitochondrial mutation spectrum in *S. conica* is unstable and has changed over time such that the “snapshot” from Duplex Sequencing of heteroplasmic SNVs does not match the average spectrum over the past millions of years in this lineage. A recent analysis of another genus with accelerated mitochondrial sequence divergence *(Ajuga)* found large increases in GC content (Liu *et al*. 2020), suggesting that bouts of accelerated and GC-biased evolution can occur in angiosperm mitochondrial genomes.

In contrast to the findings in *S. conica*, we did not observe elevated SNV frequencies in *S. noctiflora* (Figure 2) despite a comparable history of accelerated sequence evolution (Figure 1). The low SNV frequencies in *S. noctiflora* suggest that it has reverted to lower mutation rates after a past episode of acceleration. This type of reversion has been inferred based on phylogenetic evidence in other accelerated lineages such as *Plantago* and *Pelargonium* (Cho *et al*. 2004; Parkinson *et al*. 2005). We also have previously speculated that *S. noctiflora* no longer has a high mitochondrial mutation rate based on its reduced rate of sequence divergence from close relatives *S. turkestanica* and *S. undulata* and its extremely low amount of intraspecific sequence polymorphism (Sloan *et al*. 2009; Sloan *et al*. 2012a; Wu *et al*. 2015; Wu and Sloan 2019). However, if a mutation rate reversion has occurred in this lineage, it may not have simply reversed the process that led to the initial rate increase. Notably, *S. noctiflora* had a different mitochondrial mutation spectrum than the two species that have maintained low mitochondrial substitution rates throughout their histories (*A. thaliana* and *S. latifolia)*. It also retains a mitochondrial genome that is radically altered in size, structure, and apparent recombinational activity (Sloan *et al*. 2012a). Therefore, the mechanisms responsible for mitochondrial DNA replication and maintenance in *S. noctiflora* may still be quite different than in typical angiosperms despite the apparent reversion to ancestral-like rates in this species.

The above interpretations are largely based on the premise that the abundance of heteroplasmic SNVs is correlated with the mutation rate. Although it is probably a reasonable assumption that these two features are correlated, the amount of heteroplasmic genetic variation that is maintained will also depend on the (effective) population size of mitochondrial genome copies. Therefore, we cannot rule out the possibility that some of the observed differences in SNV frequency among species could be related to variation in traits such as the extent of the mtDNA transmission bottleneck during reproduction (Stewart and Chinnery 2015; Zhang *et al*. 2018; Johnston 2019). Likewise, analysis of mitochondrial SNVs in *Arabidopsis* leaf tissue has indicated that variant frequencies may be affected by features such as plant age and development (Wynn *et al*. 2020). Therefore, future studies to separate effects of mutation and population size will be useful. One possibility is that heteroplasmic SNVs identified by Duplex Sequencing could then be tracked across generations with allele-specific ddPCR. Quantifying variance in levels of inherited heteroplasmies can serve as an effective means to quantify the effective number of transmitted genome copies (Johnston 2019). However, this may be more challenging with species such as *S. latifolia* and *S. noctiflora* where inherited heteroplasmies appear to be rare.

### Mitochondrial genome copy number and recombinational repair

One unexpected finding from total-cellular shotgun sequencing was the remarkably low copy number of the mitochondrial genome is *S. conica* (Figure 3). The ratio of mitochondrial to nuclear genome copies in the analyzed leaf tissue samples implies that there is only about one to two mitochondrial genome copies per cell, under the assumption that most cells are diploid. However, this would depend on the extent of endoreduplication in *S. conica*. Species with smaller nuclear genomes have been found to undergo a greater amount of endoreduplication on average (Barow and Meister 2003), so it is possible that the ratio of organellar to nuclear genome copies is skewed downward in *S. conica* for this reason. Although there is evidence that plant cells can have far more mitochondria than mitochondrial genome copies (Preuten *et al*. 2010; Shen *et al*. 2019), it is difficult to imagine how mitochondrial function could be maintained throughout development with only one or two mitochondrial genome copies per cell. The sequencing and ddPCR datasets used to generate copynumber estimates were derived from mature rosette leaf tissue. Therefore, it is possible that this is a case of mitochondrial genome “abandonment” in tissue that is not destined for further growth or reproduction (Bendich 2013; Oldenburg *et al*. 2013; Wynn *et al*. 2020). Previous studies have suggested that some plants undergo a major decline in plastid genome copy number in mature leaves (Shaver *et al*. 2006; Rowan *et al*. 2009), although this conclusion has been the subject to substantial criticism and debate (Golczyk *et al*. 2014; Greiner *et al*. 2020). We found that all three *Silene* species retained hundreds of plastid genome copies per cell, but the dramatic differences in mitochondrial genome copy number across species have intriguing implications. An important future direction will be to characterize variation in *S. conica* mitochondrial genome copy numbers throughout development, especially in meristematic and reproductive tissues.

Even under the likely scenario that other *S. conica* tissues harbor higher mitochondrial genome copy numbers than observed in our analysis, it is possible that such values will still be unusually low compared to most plants and other eukaryotes. *Silene conica* is distinct in having one of the largest and most fragmented mitochondrial genomes ever identified (Sloan *et al*. 2012a). Such genome size and architecture might pose a challenge for mtDNA maintenance in this species. Notably, mtDNA accounted for a similar proportion of total-cellular DNA in *S. conica* and *S. latifolia* despite the ~100-fold difference in mitochondrial genome copy number between these samples because the *S. conica* mitochondrial genome is ~45-fold larger than in *S. latifolia*, and the *S. conica* nuclear genome is ~3-fold smaller than in *S. latifolia*. Nevertheless, the low mitochondrial genome copy number in *S. conica* means that any given region of the mitochondrial genome, including key functional content such as protein-coding sequence, has an unusually low stoichiometry relative to the nucleus.

We hypothesize that low mitochondrial genome copy number may be a cause of the high mutation rates in *S. conica*. It is thought that the typically low mutation rates in plant organelle genomes can be attributed to accurate DNA repair via homologous recombination (Khakhlova and Bock 2006; Christensen 2014; Ayala-García *et al*. 2018; Chevigny *et al*. 2020; Wu *et al*. 2020a). As such, the ability to maintain low rates would be sensitive to the availability of templates for recombinational repair and, thus, the number of genome copies. Notably, we did not observe elevated SNV frequencies in the *S. conica* plastid genome (Figure 2), which appears to retain a typical copy number, unlike the *S. conica* mitochondrial genome (Figure 3). This hypothesized role of copy number in recombinational repair is consistent with the observation that nucleotide substitution rates are lower in large repeats than in single-copy regions within plant organelle genomes (Wolfe *et al*. 1987; Davila *et al*. 2011; Zhu *et al*. 2016). Therefore, if the population of mitochondrial genome copies is too sparsely distributed in the cells of *S. conica*, it may be unable to make full use of recombinational repair and instead rely on less accurate forms of repair or leave some DNA damage and mismatches entirely unrepaired. In dissertation research conducted with Jeffrey Mower, Wenhu Guo (2014) arrived at a similar hypothesis after observing an unusually low mitochondrial genome copy number in *Plantago media*, another angiosperm with extremely elevated rates of mitochondrial sequence evolution (Cho *et al*. 2004). Therefore, it appears possible that correlated changes in mitochondrial mutation rate and genome copy number may have occurred many times independently in plants. Testing this hypothesized relationship between mitochondrial genome copy number and mutation rate should provide insight into the causes of extreme variation in rates of mitochondrial sequence evolution observed across angiosperms.

## ACKNOWLEDGEMENTS

We thank Justin Havird for helpful discussion and providing *S. conica* full-sib seeds and Jessica Warren for assistance with DNA extraction and figure preparation. We also thank two anonymous reviewers for insightful comments that improved the manuscript. This work was supported by a grant from the NIH (R01 GM118046) and an NSF graduate fellowship (DGE-1450032).

## SUPPLEMENTARY MATERIAL

**Figure S1.**
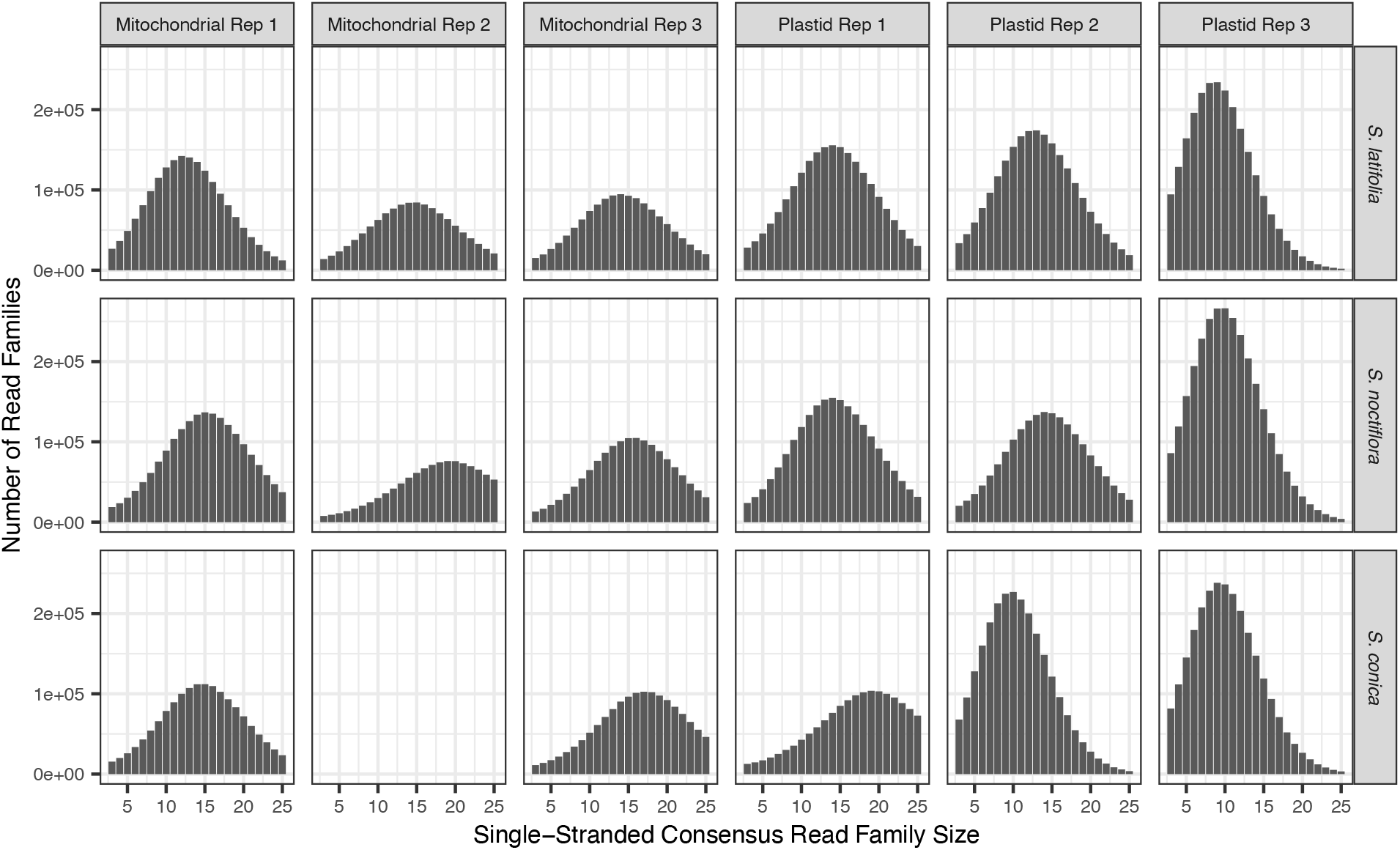
Summary of family sizes for single-stranded consensus sequences used for generation of double-stranded consensus sequences for each Duplex Sequencing library. The analysis pipeline required a minimum family size of 3 reads for each of the two complementary single-stranded families. The mitochondrial DNA library for the second biological replicate from *S. conica* was not sequenced because of an apparent library loading error.

**Figure S2.**
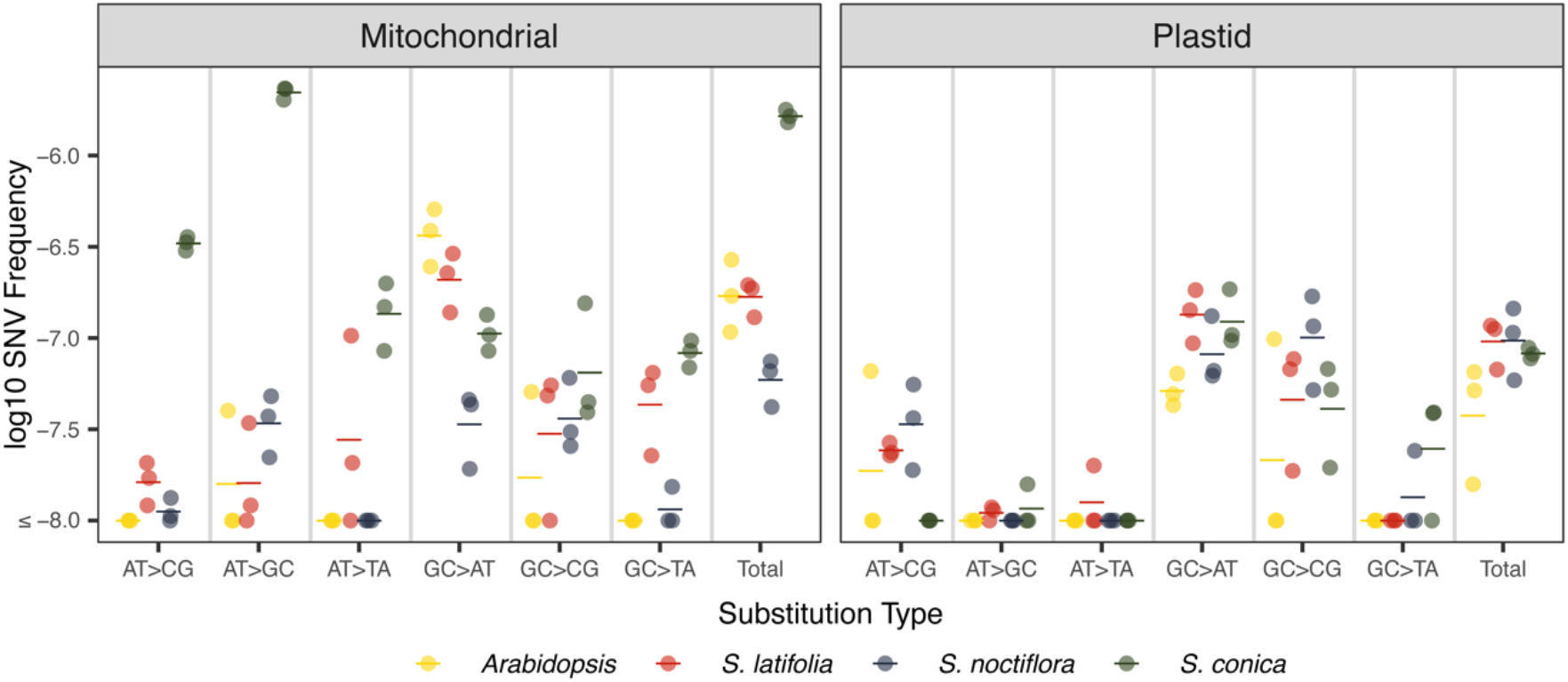
Variation in mitochondrial and plastid SNV frequencies and spectra across *Arabidopsis* and *Silene* species. The same data shown in Figure 2 are plotted on a log scale here.

**Figure S3.**
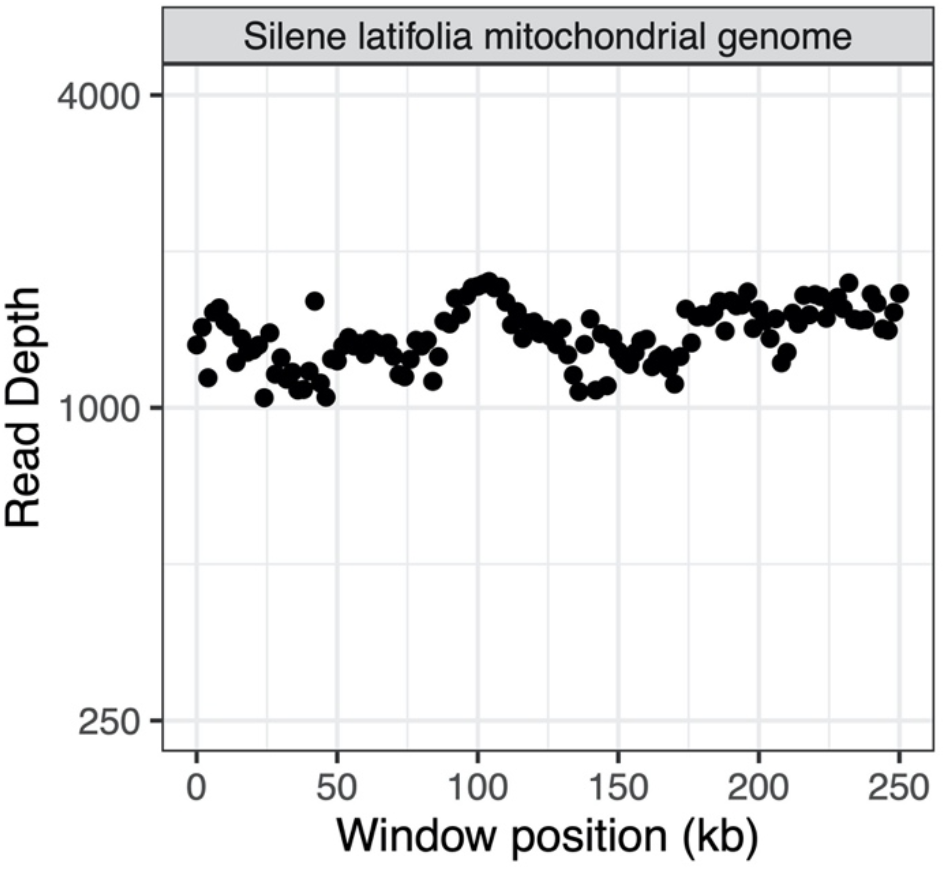
Summary of mitochondrial genome coverage in the *S. latifolia* UK2600 total-cellular sequencing library. Coverage is estimated based on reads mapping with a maximum of one mismatch and no indels in 2-kb windows.

**Figure S4.**
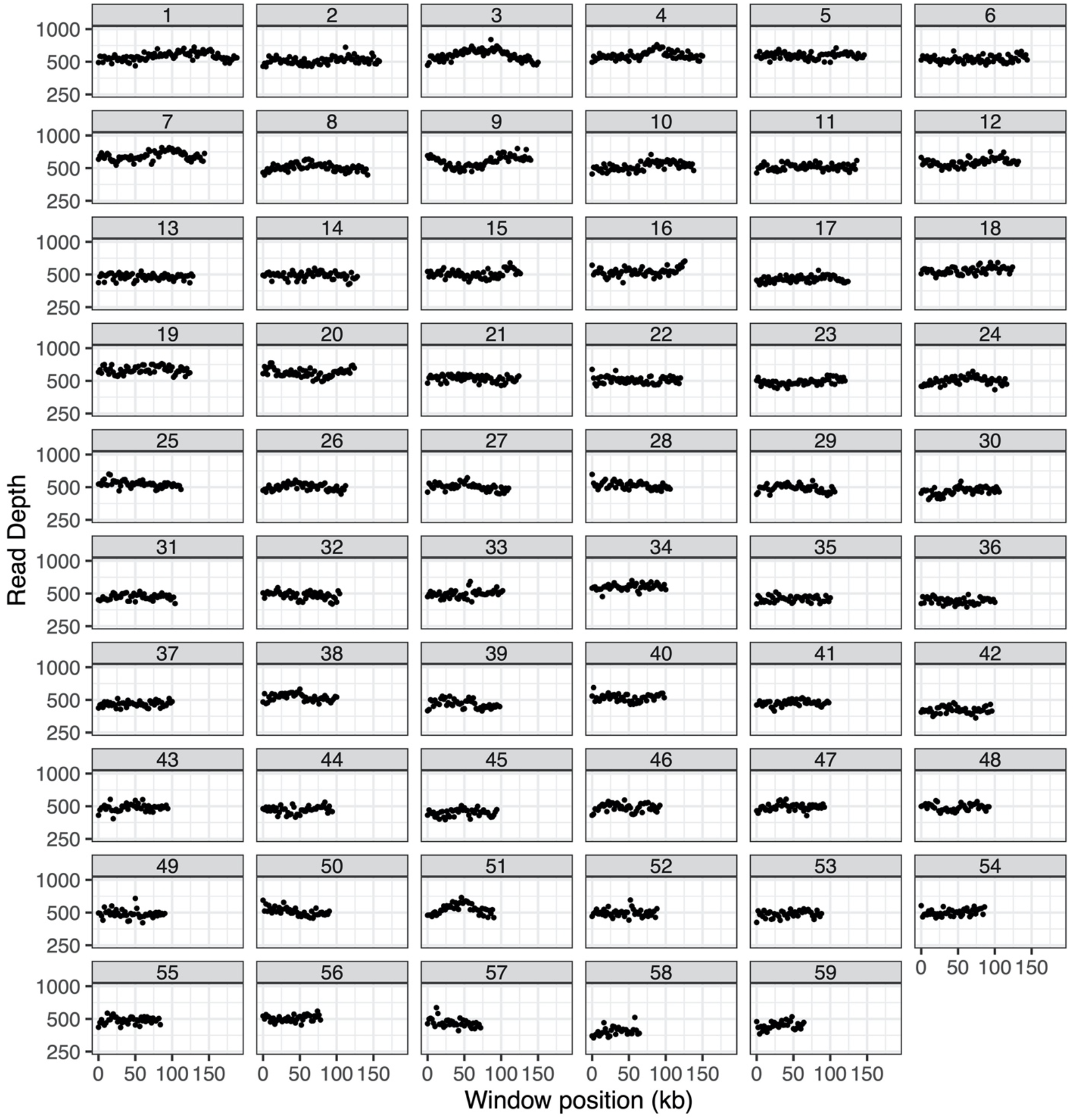
Summary of mitochondrial genome coverage in the *S. noctiflora* OSR total-cellular sequencing library. Coverage is estimated based on reads mapping with a maximum of one mismatch and no indels in 2-kb windows. Each panel represents a different chromosome within the multichromosomal genome.

**Figure S5a.**
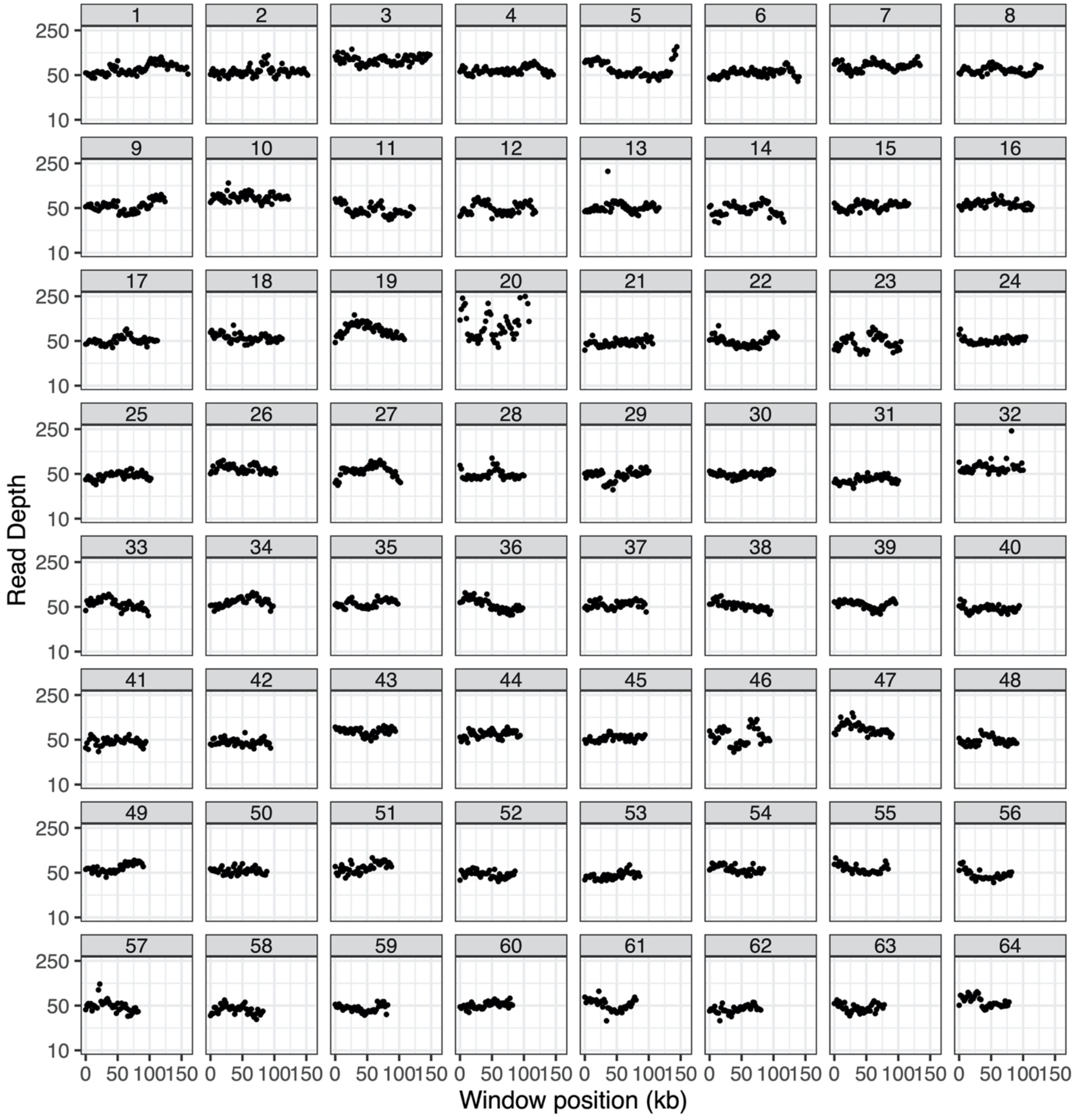
Summary of coverage for mitochondrial chromosomes 1 through 64 in the *S. conica* ABR total-cellular sequencing library. Coverage is estimated based on reads mapping with a maximum of one mismatch and no indels in 2-kb windows. Note that four of the data points on chromosome 20 and two of the data points on chromosome 32 exceeded a coverage of 250× and are not shown to improve readability of the plots.

**Figure S5b.**
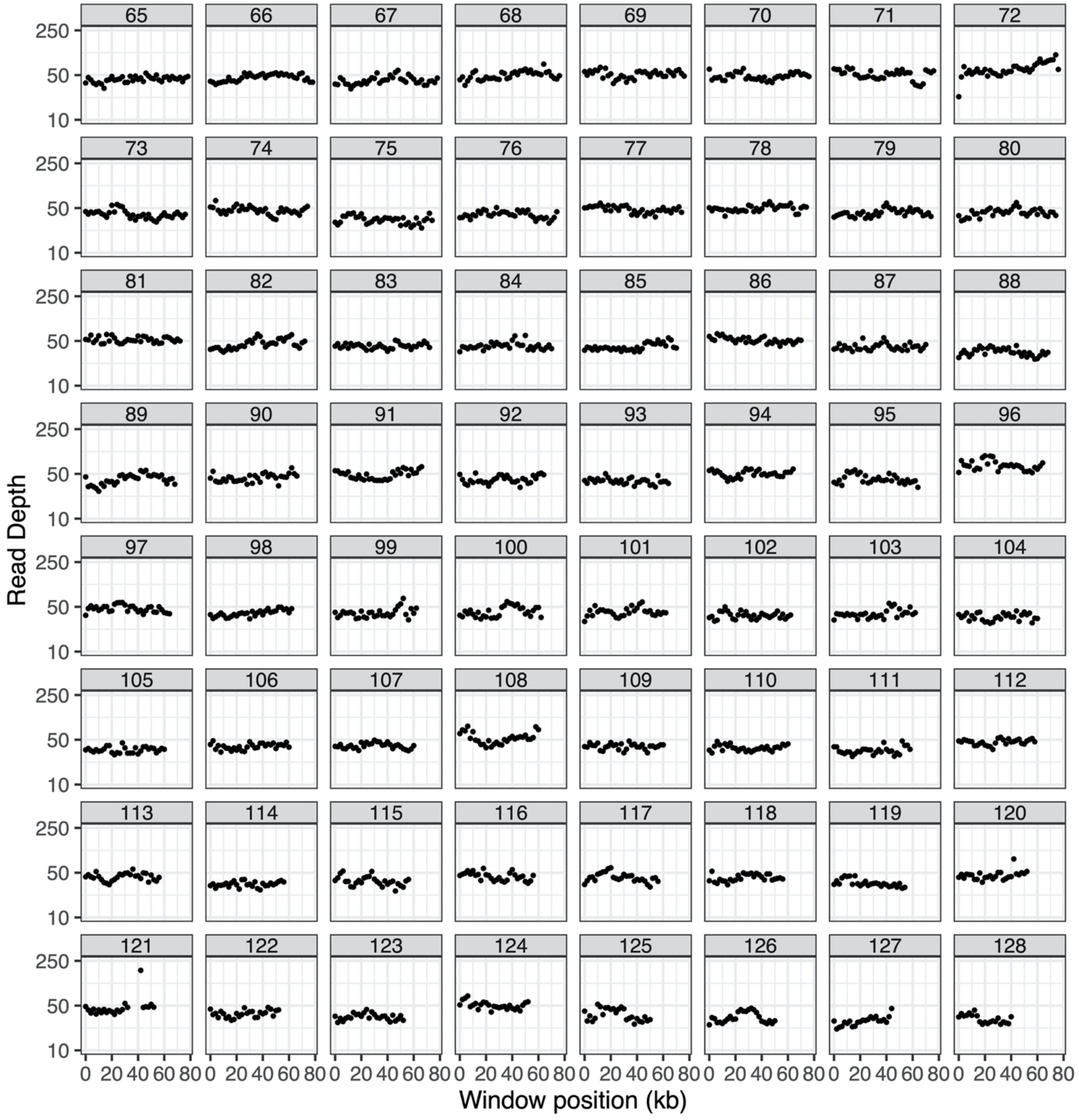
Summary of coverage for mitochondrial chromosomes 65 through 128 in the *S. conica* ABR total-cellular sequencing library. Coverage is estimated based on reads mapping with a maximum of one mismatch and no indels in 2-kb windows. Note that four of the data points on chromosome 121 exceeded a coverage of 250× and are not shown to improve readability of the plots.

**Table S1.**
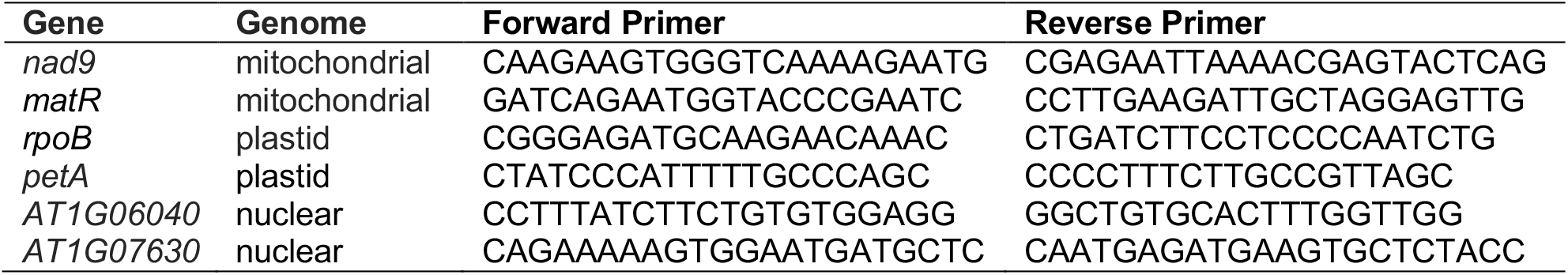
Primers used in the *S. conica* ddPCR analysis. The two nuclear markers are referred to by the identifiers for the homologs in *Arabidopsis thaliana*.

**Table S2.** Summary of library sequencing and mapping. Reported coverage values and mapping percentages are for duplex consensus sequences.

| Library | Library Type | SRA Accession | Raw Read Pairs | Mito Coverage (bp) | Plastid Coverage (bp) | Mito Mapping (%) | Plastid Mapping (%) | Unmapped (%) |
| --- | --- | --- | --- | --- | --- | --- | --- | --- |
| Silene conica chloroplast 1 | Chloroplast duplex | SAMN17011097 | 1.15E+08 | 9.25E+07 | 2.03E+08 | 29.49% | 64.88% | 5.63% |
| Silene conica chloroplast 2 | Chloroplast duplex | SAMN17011098 | 1.15E+08 | 1.33E+08 | 2.72E+08 | 31.48% | 64.13% | 4.39% |
| Silene conica chloroplast 3 | Chloroplast duplex | SAMN17011099 | 1.42E+08 | 1.37E+08 | 2.70E+08 | 31.72% | 62.55% | 5.73% |
| Silene conica mito 1 | Mitochondrial duplex | SAMN17011100 | 1.01E+08 | 2.18E+08 | 1.77E+07 | 80.87% | 6.58% | 12.55% |
| Silene conica mito 3 | Mitochondrial duplex | SAMN17011101 | 1.14E+08 | 2.16E+08 | 2.01E+07 | 77.21% | 7.16% | 15.63% |
| Silene latifolia chloroplast 1 | Chloroplast duplex | SAMN17011085 | 1.33E+08 | 2.89E+07 | 2.83E+08 | 7.39% | 72.35% | 20.26% |
| Silene latifolia chloroplast 2 | Chloroplast duplex | SAMN17011086 | 1.28E+08 | 3.54E+07 | 2.41E+08 | 8.82% | 60.10% | 31.08% |
| Silene latifolia chloroplast 3 | Chloroplast duplex | SAMN17011087 | 1.01E+08 | 3.68E+07 | 2.73E+08 | 9.13% | 67.73% | 23.14% |
| Silene latifolia mito 1 | Mitochondrial duplex | SAMN17011088 | 1.13E+08 | 1.16E+08 | 2.67E+07 | 38.26% | 8.83% | 52.91% |
| Silene latifolia mito 2 | Mitochondrial duplex | SAMN17011089 | 9.79E+07 | 4.93E+07 | 1.63E+07 | 24.87% | 8.21% | 66.92% |
| Silene latifolia mito 3 | Mitochondrial duplex | SAMN17011090 | 8.83E+07 | 6.56E+07 | 2.23E+07 | 27.94% | 9.48% | 62.58% |
| Silene noctiflora chloroplast 1 | Chloroplast duplex | SAMN17011091 | 1.37E+08 | 1.06E+08 | 2.55E+08 | 27.63% | 66.35% | 6.02% |
| Silene noctiflora chloroplast 2 | Chloroplast duplex | SAMN17011092 | 1.25E+08 | 9.39E+07 | 2.20E+08 | 28.18% | 66.12% | 5.70% |
| Silene noctiflora chloroplast 3 | Chloroplast duplex | SAMN17011093 | 1.37E+08 | 1.25E+08 | 3.19E+08 | 25.69% | 65.36% | 8.95% |
| Silene noctiflora mito 1 | Mitochondrial duplex | SAMN17011094 | 1.32E+08 | 2.76E+08 | 1.56E+07 | 83.03% | 4.68% | 12.29% |
| Silene noctiflora mito 2 | Mitochondrial duplex | SAMN17011095 | 1.06E+08 | 1.88E+08 | 1.22E+07 | 81.54% | 5.28% | 13.18% |
| Silene noctiflora mito 3 | Mitochondrial duplex | SAMN17011096 | 1.03E+08 | 1.93E+08 | 2.39E+07 | 71.53% | 8.84% | 19.63% |
| Silene conica ABR | Total cellular shotgun | SAMN17011105 | 4.98E+08 | 3.53E+08 | 1.68E+09 | 0.49% | 2.31% | 97.20% |
| Silene noctiflora OSR | Total cellular shotgun | SAMN17011104 | 5.76E+08 | 3.45E+09 | 2.10E+09 | 2.51% | 1.53% | 95.96% |
| Silene latifolia UK2600 | Total cellular shotgun | SAMN17011102 | 2.81E+08 | 5.95E+08 | 5.51E+09 | 0.50% | 4.64% | 94.86% |
| Silene latifolia Kew 32982 | Total cellular shotgun | SAMN17011103 | 3.04E+08 |  |  |  |  |  |

**Table S3.** *Silene* mitochondrial SNV count and frequency data with and without *k*-mer filtering

| Species | Rep | k-mer Filt | SNV Counts |  |  |  |  |  |  | SNV Frequencies |  |  |  |  |  |  |
| --- | --- | --- | --- | --- | --- | --- | --- | --- | --- | --- | --- | --- | --- | --- | --- | --- |
|  |  |  | A>C | A>G | A>T | C>A | C>G | C>T | Total | A>C | A>G | A>T | C>A | C>G | C>T | Total |
| *Silene conica* | 1 | 10 | 53 | 356 | 26 | 13 | 6 | 18 | 472 | 3.0E-07 | 2.0E-06 | 1.5E-07 | 9.7E-08 | 4.5E-08 | 1.3E-07 | 1.5E-06 |
| *Silene conica* | 2 | 10 | 27 | 176 | 15 | 4 | 9 | 6 | 237 | 3.6E-07 | 2.3E-06 | 2.0E-07 | 6.9E-08 | 1.6E-07 | 1.0E-07 | 1.8E-06 |
| *Silene conica* | 3 | 10 | 67 | 462 | 17 | 13 | 6 | 13 | 578 | 3.3E-07 | 2.3E-06 | 8.5E-08 | 8.5E-08 | 3.9E-08 | 8.5E-08 | 1.6E-06 |
| *Silene conica* | 1 | None | 150 | 483 | 58 | 49 | 12 | 217 | 969 | 8.5E-07 | 2.7E-06 | 3.3E-07 | 3.6E-07 | 8.9E-08 | 1.6E-06 | 3.1E-06 |
| *Silene conica* | 2 | None | 64 | 238 | 24 | 20 | 12 | 98 | 456 | 8.5E-07 | 3.2E-06 | 3.2E-07 | 3.5E-07 | 2.1E-07 | 1.7E-06 | 3.4E-06 |
| *Silene conica* | 3 | None | 158 | 610 | 50 | 59 | 17 | 231 | 1125 | 7.9E-07 | 3.0E-06 | 2.5E-07 | 3.9E-07 | 1.1E-07 | 1.5E-06 | 3.2E-06 |
| *Silene latifolia* | 1 | 10 | 1 | 1 | 0 | 4 | 3 | 18 | 27 | 5.7E-09 | 5.7E-09 | 0.0E+00 | 3.0E-08 | 2.2E-08 | 1.3E-07 | 8.7E-08 |
| *Silene latifolia* | 2 | 10 | 1 | 0 | 1 | 2 | 2 | 5 | 11 | 1.3E-08 | 0.0E+00 | 1.3E-08 | 3.5E-08 | 3.5E-08 | 8.6E-08 | 8.2E-08 |
| *Silene latifolia* | 3 | 10 | 1 | 2 | 6 | 1 | 0 | 10 | 20 | 5.0E-09 | 1.0E-08 | 3.0E-08 | 6.5E-09 | 0.0E+00 | 6.5E-08 | 5.7E-08 |
| *Silene latifolia* | 1 | None | 9 | 8 | 6 | 17 | 5 | 42 | 87 | 1.1E-07 | 9.7E-08 | 7.3E-08 | 2.7E-07 | 8.1E-08 | 6.8E-07 | 6.0E-07 |
| *Silene latifolia* | 2 | None | 5 | 8 | 14 | 17 | 6 | 58 | 108 | 1.0E-07 | 1.7E-07 | 2.9E-07 | 4.7E-07 | 1.7E-07 | 1.6E-06 | 1.3E-06 |
| *Silene latifolia* | 3 | None | 9 | 16 | 18 | 19 | 10 | 52 | 124 | 1.5E-07 | 2.7E-07 | 3.1E-07 | 4.3E-07 | 2.3E-07 | 1.2E-06 | 1.2E-06 |
| *Silene noctiflora* | 1 | 10 | 3 | 5 | 1 | 0 | 4 | 3 | 16 | 1.3E-08 | 2.2E-08 | 4.4E-09 | 0.0E+00 | 2.6E-08 | 1.9E-08 | 4.2E-08 |
| *Silene noctiflora* | 2 | 10 | 1 | 8 | 0 | 0 | 7 | 5 | 21 | 6.0E-09 | 4.8E-08 | 0.0E+00 | 0.0E+00 | 6.1E-08 | 4.3E-08 | 7.4E-08 |
| *Silene noctiflora* | 3 | 10 | 2 | 7 | 0 | 2 | 4 | 6 | 21 | 1.1E-08 | 3.7E-08 | 0.0E+00 | 1.5E-08 | 3.1E-08 | 4.6E-08 | 6.6E-08 |
| *Silene noctiflora* | 1 | None | 3 | 8 | 3 | 4 | 6 | 12 | 36 | 1.3E-08 | 3.5E-08 | 1.3E-08 | 2.6E-08 | 3.8E-08 | 7.7E-08 | 9.4E-08 |
| *Silene noctiflora* | 2 | None | 2 | 11 | 1 | 2 | 13 | 9 | 38 | 1.2E-08 | 6.6E-08 | 6.0E-09 | 1.7E-08 | 1.1E-07 | 7.8E-08 | 1.3E-07 |
| *Silene noctiflora* | 3 | None | 5 | 13 | 2 | 11 | 9 | 11 | 51 | 2.7E-08 | 6.9E-08 | 1.1E-08 | 8.4E-08 | 6.9E-08 | 8.4E-08 | 1.6E-07 |

**Table S4.** *Silene* plastid SNV count and frequency data with and without *k*-mer filtering

| Species | Rep | k-mer Filt | SNV Counts |  |  |  |  |  |  | SNV Frequencies |  |  |  |  |  |  |
| --- | --- | --- | --- | --- | --- | --- | --- | --- | --- | --- | --- | --- | --- | --- | --- | --- |
|  |  |  | A>C | A>G | A>T | C>A | C>G | C>T | Total | A>C | A>G | A>T | C>A | C>G | C>T | Total |
| *Silene conica* | 1 | 10 | 0 | 2 | 1 | 3 | 4 | 8 | 18 | 0.0E+00 | 1.6E-08 | 7.9E-09 | 3.9E-08 | 5.2E-08 | 1.0E-07 | 8.8E-08 |
| *Silene conica* | 2 | 10 | 0 | 0 | 0 | 4 | 7 | 10 | 21 | 0.0E+00 | 0.0E+00 | 0.0E+00 | 3.9E-08 | 6.8E-08 | 9.7E-08 | 7.7E-08 |
| *Silene conica* | 3 | 10 | 0 | 1 | 0 | 0 | 2 | 19 | 22 | 0.0E+00 | 6.0E-09 | 0.0E+00 | 0.0E+00 | 2.0E-08 | 1.9E-07 | 8.2E-08 |
| *Silene conica* | 1 | None | 1 | 4 | 1 | 3 | 4 | 12 | 25 | 7.9E-09 | 3.2E-08 | 7.9E-09 | 3.9E-08 | 5.2E-08 | 1.6E-07 | 1.2E-07 |
| *Silene conica* | 2 | None | 0 | 0 | 0 | 6 | 8 | 20 | 34 | 0.0E+00 | 0.0E+00 | 0.0E+00 | 5.8E-08 | 7.7E-08 | 1.9E-07 | 1.3E-07 |
| *Silene conica* | 3 | None | 0 | 1 | 1 | 2 | 4 | 26 | 34 | 0.0E+00 | 6.0E-09 | 6.0E-09 | 2.0E-08 | 3.9E-08 | 2.5E-07 | 1.3E-07 |
| *Silene latifolia* | 1 | 10 | 4 | 2 | 0 | 1 | 2 | 10 | 19 | 2.3E-08 | 1.1E-08 | 0.0E+00 | 9.4E-09 | 1.9E-08 | 9.4E-08 | 6.7E-08 |
| *Silene latifolia* | 2 | 10 | 4 | 0 | 3 | 0 | 7 | 13 | 27 | 2.7E-08 | 0.0E+00 | 2.0E-08 | 0.0E+00 | 7.7E-08 | 1.4E-07 | 1.1E-07 |
| *Silene latifolia* | 3 | 10 | 4 | 2 | 0 | 0 | 7 | 19 | 32 | 2.4E-08 | 1.2E-08 | 0.0E+00 | 0.0E+00 | 6.7E-08 | 1.8E-07 | 1.2E-07 |
| *Silene latifolia* | 1 | None | 8 | 6 | 4 | 2 | 3 | 31 | 54 | 4.5E-08 | 3.4E-08 | 2.3E-08 | 1.9E-08 | 2.8E-08 | 2.9E-07 | 1.9E-07 |
| *Silene latifolia* | 2 | None | 10 | 4 | 11 | 10 | 10 | 38 | 83 | 6.7E-08 | 2.7E-08 | 7.3E-08 | 1.1E-07 | 1.1E-07 | 4.2E-07 | 3.4E-07 |
| *Silene latifolia* | 3 | None | 7 | 12 | 6 | 6 | 9 | 30 | 70 | 4.1E-08 | 7.1E-08 | 3.5E-08 | 5.8E-08 | 8.7E-08 | 2.9E-07 | 2.6E-07 |
| *Silene noctiflora* | 1 | 10 | 3 | 0 | 1 | 0 | 5 | 6 | 15 | 1.9E-08 | 0.0E+00 | 6.3E-09 | 0.0E+00 | 5.2E-08 | 6.2E-08 | 5.9E-08 |
| *Silene noctiflora* | 2 | 10 | 5 | 0 | 0 | 2 | 14 | 11 | 32 | 3.6E-08 | 0.0E+00 | 0.0E+00 | 2.4E-08 | 1.7E-07 | 1.3E-07 | 1.5E-07 |
| *Silene noctiflora* | 3 | 10 | 11 | 0 | 1 | 0 | 14 | 8 | 34 | 5.6E-08 | 0.0E+00 | 5.1E-09 | 0.0E+00 | 1.2E-07 | 6.6E-08 | 1.1E-07 |
| *Silene noctiflora* | 1 | None | 5 | 1 | 1 | 1 | 5 | 9 | 22 | 3.1E-08 | 6.3E-09 | 6.3E-09 | 1.0E-08 | 5.2E-08 | 9.3E-08 | 8.6E-08 |
| *Silene noctiflora* | 2 | None | 6 | 1 | 0 | 2 | 14 | 19 | 42 | 4.4E-08 | 7.3E-09 | 0.0E+00 | 2.4E-08 | 1.7E-07 | 2.3E-07 | 1.9E-07 |
| *Silene noctiflora* | 3 | None | 12 | 0 | 2 | 2 | 14 | 17 | 47 | 6.1E-08 | 0.0E+00 | 1.0E-08 | 1.7E-08 | 1.2E-07 | 1.4E-07 | 1.5E-07 |

**Table S5.**
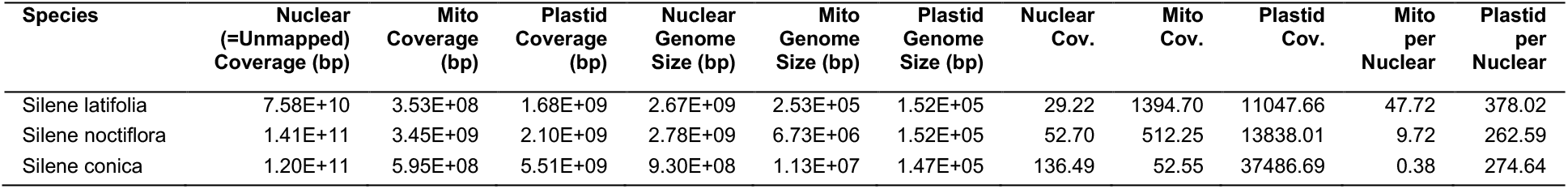
Genome copy number estimates

**Table S6.** ddPCR droplet counts and copy number calculations

| Sample | Genome | Marker | Positive droplets | Negative droplets | Copies per reaction | Dilution Factor | Copies per nuclear copy |
| --- | --- | --- | --- | --- | --- | --- | --- |
| New1 | Mito | matR | 1244 | 6955 | 3967 | 1 | 0.98 |
| New2 | Mito | matR | 2413 | 10431 | 4900 | 1 | 1.09 |
| New3 | Mito | matR | 1987 | 11250 | 3820 | 1 | 0.91 |
| Original | Mito | matR | 2356 | 13893 | 3680 | 1 | 0.42 |
| New1 | Mito | nad9 | 1842 | 10249 | 3880 | 1 | 0.96 |
| New2 | Mito | nad9 | 1202 | 4781 | 5280 | 1 | 1.18 |
| New3 | Mito | nad9 | 990 | 5106 | 4160 | 1 | 1.00 |
| Original | Mito | nad9 | 2268 | 13951 | 3540 | 1 | 0.41 |
| New1 | Plastid | petA | 6502 | 7514 | 14660 | 200 | 727.54 |
| New2 | Plastid | petA | 6692 | 6199 | 17220 | 200 | 767.04 |
| New3 | Plastid | petA | 6392 | 8264 | 13480 | 200 | 644.98 |
| Original | Plastid | petA | 5601 | 1651 | 34820 | 200 | 804.16 |
| New1 | Plastid | rpoB | 5868 | 6456 | 15220 | 200 | 755.33 |
| New2 | Plastid | rpoB | 6803 | 6015 | 17800 | 200 | 792.87 |
| New3 | Plastid | rpoB | 6409 | 8078 | 13740 | 200 | 657.42 |
| Original | Plastid | rpoB | 10798 | 4448 | 28980 | 200 | 669.28 |
| New1 | Nuclear | AT1G06040 | 1524 | 7912 | 4140 | 1 |  |
| New2 | Nuclear | AT1G06040 | 1751 | 8889 | 4240 | 1 |  |
| New3 | Nuclear | AT1G06040 | 2070 | 10630 | 4180 | 1 |  |
| Original | Nuclear | AT1G06040 | 3982 | 10165 | 7780 | 1 |  |
| New1 | Nuclear | AT1G07630 | 1756 | 9820 | 3920 | 1 |  |
| New2 | Nuclear | AT1G07630 | 2242 | 10039 | 4740 | 1 |  |
| New3 | Nuclear | AT1G07630 | 2118 | 10890 | 4180 | 1 |  |
| Original | Nuclear | AT1G07630 | 5315 | 10629 | 9540 | 1 |  |

**Table S7.**
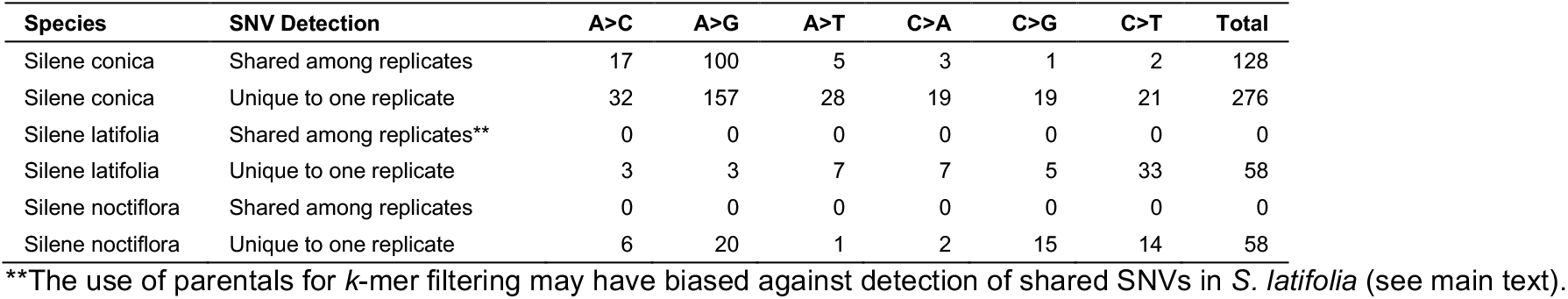
Detection of mitochondrial SNVs in multiple biological replicates

| Species | SNV Detection | A>C | A>G | A>T | C>A | C>G | C>T | Total |
| --- | --- | --- | --- | --- | --- | --- | --- | --- |
| Silene conica | Shared among replicates | 17 | 100 | 5 | 3 | 1 | 2 | 128 |
| Silene conica | Unique to one replicate | 32 | 157 | 28 | 19 | 19 | 21 | 276 |
| Silene latifolia | Shared among replicates** | 0 | 0 | 0 | 0 | 0 | 0 | 0 |
| Silene latifolia | Unique to one replicate | 3 | 3 | 7 | 7 | 5 | 33 | 58 |
| Silene noctiflora | Shared among replicates | 0 | 0 | 0 | 0 | 0 | 0 | 0 |
| Silene noctiflora | Unique to one replicate | 6 | 20 | 1 | 2 | 15 | 14 | 58 |

\*\*The use of parentals for *k*-mer filtering may have biased against detection of shared SNVs in *S. latifolia* (see main text).

